# Genopyc: a python library for investigating the genomic basis of complex diseases

**DOI:** 10.1101/2024.01.11.575316

**Authors:** Francesco Gualdi, Baldomero Oliva, Janet Piñero

## Abstract

**Motivation:** Understanding the genetic basis of complex diseases is a paramount challenge in modern genomics. However, current tools often lack the versatility to efficiently analyze the intricate relation-ships between genetic variations and disease outcomes. To address this, we introduce Genopyc, a novel Python library designed for comprehensive investigation of the genetics underlying complex dis-eases. Genopyc offers an extensive suite of functions for heterogeneous data mining and visualization, enabling researchers to delve into and integrate biological information from large-scale genomic da-tasets with ease.

**Results:** In this study, we present the Genopyc library through application to real-world genome wide association studies variants. Using Genopyc to investigate variants associated to intervertebral disc degeneration (IDD) enabled a deeper understanding of the potential dysregulated pathways involved in the disease, which can be explored and visualized by exploiting the functionalities featured in the package. Genopyc emerges as a powerful asset for researchers, fostering advancements in the un-derstanding of complex diseases and thus paving the way for more targeted therapeutic interventions. Availability: Genopyc is available at pip (https://pypi.org/project/genopyc/) and the source code of Genopyc is available at https://github.com/freh-g/genopyc

**Supplementary information:** supplementary data are available at *Bioinformatics* online.

## 1 Introduction

The onset of complex disorders is influenced by a multitude of components that include lifestyle, diet, environmental and genetic factors. In the last decades genome wide association studies (GWAS) have emerged as a powerful tool to investigate the genetic architecture underlying complex diseases (Bush 2012). However, now that we have successfully discovered thousands of genetic risk factors for numerous phenotypes, we are facing another challenge: the interpretation of these associations in the biological context; we are thus entering in the so called post-GWAS Era (Gallagher 2018). Understanding how genetic variants are translated into biological pathways remains a challenging task (Edwards 2013) and led to the development of numerous approaches to interpret GWAS results (see (Uffelmann 2021) for a comprehensive review of the type of analysis and tools).

**Figure 1.**
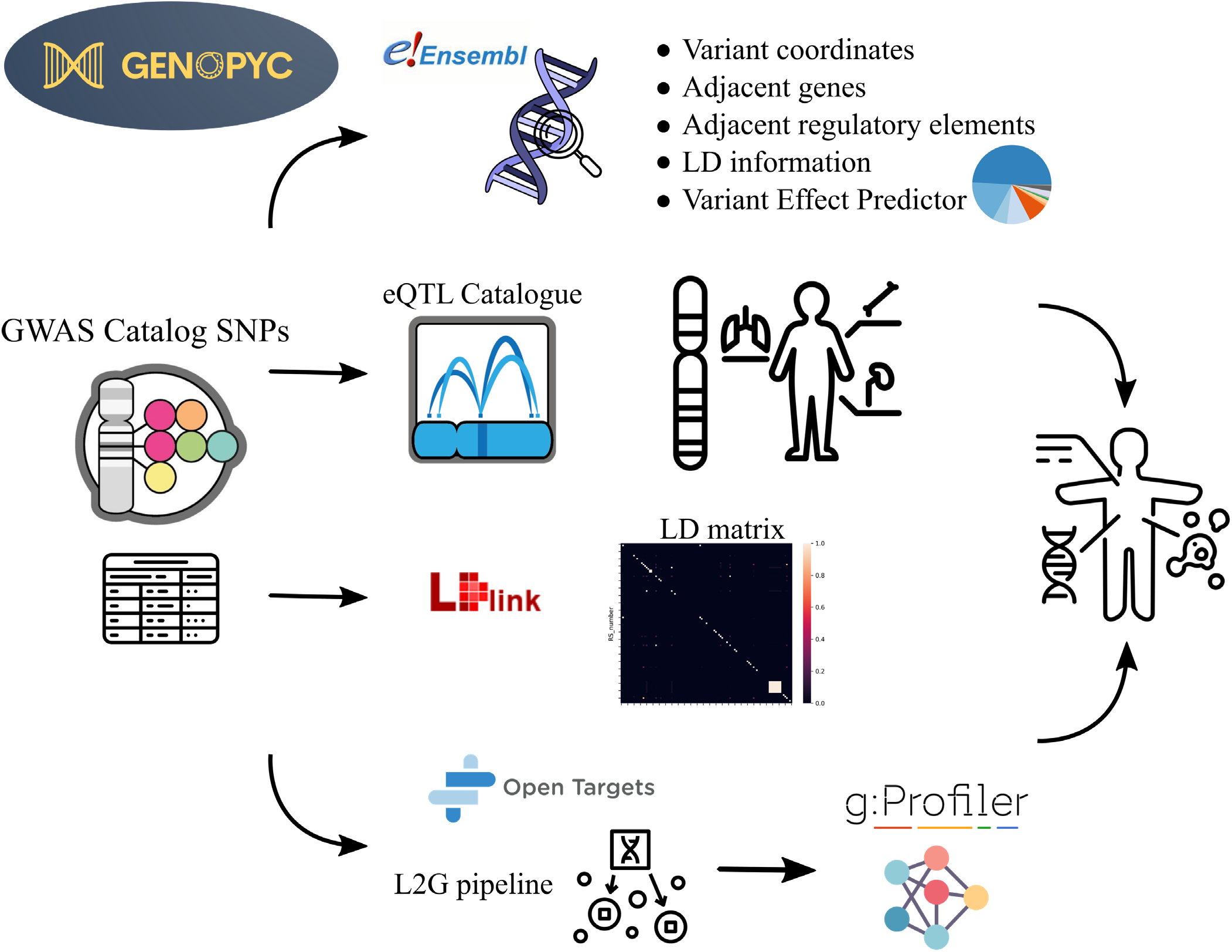
Schematic representation of the main Genopyc features and knowledge bases that are accessed.

Ascertaining the precise functional implications of the genetic variants discovered through GWAS has proven to be a formidable challenge due to the extensive amount of data required to perform these studies (Edwards 2013). In response, a plethora of novel methodological approaches has emerged to address this knowledge gap (Mulder 2017). These techniques rely on the large-scale omics datasets and repositories available to researchers such as Gene Expression Omnibus (Edgar 2002), the genotype – tissue expression project (Lonsdale 2013) and the Encode project (de Souza 2012). The enormous amount of data regarding genes and variants associated to diseases is collected in knowledge bases such as the GWAS Catalog (Sollis 2023), and DisGeNET (Piñero 2019) that offers a standardized integration from different sources. However, 90% of the genetic variation associated to complex diseases are non-coding type and a benchmark of methods to interpret how they alter genes, perturb biological pathways and ultimately lead to disease is still missing (Li 2021). Moreover, the application and integration of different tools to analyze GWAS data lead to discordant results, thus an unbiased assessment of the methods available is still required (Pérez-Granado 2022). An advancement in associating genes to non-coding variants has been made by the Open Target Genetics platform, which implemented a pipeline consisting of a machine learning model that uses heterogeneous features such as distance from variant to the gene, expression quantitative trait loci, chromatin conformation and variant effect predictor. This method outperformed the naïve distance-based methods in the prioritization of causal genes related to complex diseases loci (Mountjoy 2021). Finally, the tools and data repositories useful for analyzing this type of data are scattered, which makes understanding GWAS results a time-consuming process.

In this context we present Genopyc, a python library for investigating genetic basis of complex diseases.

Genopyc allows users to programmatically access multiple sources with the aim of understanding how non-coding variants could impact the biological pathways and thus infer the mechanisms underlying the development of complex diseases. Moreover, being fully integrated in python, all the downstream statistical analysis can be carried out in the same environment.

## 1 Implementation and features

Genopyc is a python package integrating information from several knowledgebases. The tool can receive as an input a trait, coded with Experimental Factor Ontology identifiers (Malone 2010), or the results of a GWA study. If an EFO code is used as an input, the variants associated to the trait are retrieved from the GWAS Catalog. Information such as the β coefficient, standard error, risk allele frequency and the mapped genes related to the GWAS is also retrieved. Additionally, other features such as genomic coordinates, functions whose perturbation could ultimately lead to the disease. Genopyc package also offers a functionality to visualize the results of the functional enrichment as an interactive network (see Supplementary material). In this network, genes of interest are mapped to a protein-protein interaction network derived from the HIPPIE database (Alanis-Lobato 2017) in which nodes are used to represent the gene products and edges correspond to the physical interactions between proteins. A dropdown menu allows the user to select the function enriched in the gene set and, when a function is selected, the gene-products belonging to that function are highlighted.

Genopyc can also retrieve a linkage-disequilibrium (LD) matrix for a set of SNPs by using LDlink (Machiela 2015), converting genome coordinates between genome versions and retrieving genes coordinates in the genome. LDlink calculates the LD matrix through the population-specific 1000 genomes haplotype panels (Auton 2015). Retrieving genome coordinates and mapping between genome builds are made possible by accessing Ensembl genome browser. A comparison between the main functionalities of Genopyc linkage disequilibrium (LD) correlated SNPs and neighboring functional elements can be obtained from Ensembl Genome Browser (Martin 2023).

Once the variants are retrieved, Genopyc queries the variant effect predictor (VEP) to predict the consequences of the SNPs on the transcript and its effect on neighboring genes and functional elements (McLaren 2016). Often SNPs associated to complex phenotypes fall in non-coding regions of the genome and are more likely to have regulatory effects (Prokunina 2004). Therefore, it is possible to retrieve the expression quantitative trait loci (eQTL) related to variants through the eQTL Catalogue (Kerimov 2021). Finally, Genopyc integrates the locus to gene (L2G) pipeline from Open Target Genetics to uncover the target gene or genes of variants located in non-coding regions. Once a variant is associated to a gene or genes, the significantly enriched pathways are retrieved through G:Profiler (Raudvere 2019). In this way the user can elucidate the and other tools for post-GWAS analysis is shown in Table 1. Genopyic is the only library that integrates multiple analysis to connect variants to genes (conditional, colocalization, fine mapping), gather functional information to annotate variants (eQTLs, HI-C, linkage disequlibrium, VEP, functional genomic elements) and perform functional enrichment to detect possible pathways perturbed by genetic variations. In summary we provide an all-in-one tool to retrieve and interpret the effect of genomic variants on the development of complex disease. Genopyc is easily installable via pip and can be integrated into python environments being built upon main python libraries.

**Table 1.** Comparison between Genopyc and the main tools for post GWAS analysis. Genopyc integrates diverse functionalities allowing a more flexible investigation of variants related to diseases

| Tool | Retrieve trait associated variants | Conditional analysis | Fine mapping | eQTL Co-localization | HI-C | Functional Enrichment | Linkage Disequilibrium | Genomic features | Variant annotation |
| --- | --- | --- | --- | --- | --- | --- | --- | --- | --- |
| Genopyc | ✓ | ✓ | ✓ | ✓ | ✓ | ✓ | ✓ | ✓ | ✓ |
| Coloc | × | × | × | ✓ | × | × | × | × | ✓ |
| FUMA | × | ✓ | × | ✓ | ✓ | × | ✓ | × | × |
| Finemap | × | × | ✓ | × | × | × | ✓ | × | × |
| Ensemble API | × | × | × | × | × | × | ✓ | ✓ | ✓ |
| Open Targets: Genetics | ✓ | ✓ | ✓ | ✓ | ✓ | × | ✓ | × | ✓ |

## 2 Use Case

To illustrate the utility of Genopyc, we applied it to the variants associated to lumbar disc degeneration that are available in the GWAS catalog (IDD, EFO:0004994). IDD is a complex multifactorial condition for which the onset and its molecular mechanisms are poorly understood. Thanks to Genopyc and multiple data integration we highlight the involvement of variants associated downstream to pathways that may be relevant to the IDD, such as SP1 (Xu 2016), HIF1-α (Meng 2018) and AP-2α (Li 2020), that according to the literature are tightly associated with IDD. Conversely, the functional enrichment didn’t bring any result or valuable information on the pathways that could be dysregulated in the disease. This example highlights that thanks to Genopyc a user can unveil a greater understanding of human complex traits.

## 3 Availability

The library can be installed via pip: https://pypi.org/project/genopyc/ The source code is available at: https://github.com/freh-g/genopyc The notebook with the use case is available at: https://github.com/freh-g/genopyc

## Supporting information

Function enrichment visualization tool

## References

Alanis-Lobato, Gregorio, Andrade-Navarro Miguel A., Schaefer Martin H. HIPPIE v2.0: Enhancing meaningfulness and reliability of proteinprotein interaction networks Nucleic Acids Res 2017;45:D408–D414.

Auton, Adam et al. A global reference for human genetic variation Nature 2015;526:68–74.

Bovonratwet, Patawut et al. Identification of Novel Genetic Markers for the Risk of Spinal Pathologies: A Genome-Wide Association Study of 2 Biobanks JBJS 2023;105

Bush, William S., Moore Jason H. Chapter 11: Genome-Wide Association Studies PLoS Comput Biol 2012;8

de Souza, Natalie Genomics: The ENCODE project Nat Methods 2012

Edgar, Ron, Domrachev, Michael, Lash Alex E Gene Expression Omnibus: NCBI gene expression and hybridization array data repository 2002;30

Edwards, Stacey L. et al. Beyond GWASs: Illuminating the dark road from association to function Am J Hum Genet 2013;93:779–797.

Gallagher, Michael D., Chen-Plotkin Alice S. The Post-GWAS Era: From Association to Function Am J Hum Genet 2018;102:717–730.

Kerimov, Nurlan et al. A compendium of uniformly processed human gene expression and splicing quantitative trait loci Nat Genet 2021;53:1290–1299.

Li, Binglan, Ritchie Marylyn D. From GWAS to Gene: Transcriptome-Wide Association Studies and Other Methods to Functionally Understand GWAS Discoveries Front Genet 2021;12

Li, Haoxi et al. Role of AP-2α/TGF-ß1/Smad3 axis in rats with intervertebral disc degeneration Life Sci 2020;263

Lonsdale, John et al. The Genotype-Tissue Expression (GTEx) project Nat Genet 2013;45:580–585.

Machiela, Mitchell J., Chanock Stephen J. LDlink: A web-based application for exploring population-specific haplotype structure and linking correlated alleles of possible functional variants Bioinformatics 2015;31:3555–3557.

Malone, James et al. Modeling sample variables with an Experimental Factor Ontology Bioinformatics 2010;26:1112–1118.

Martin, Fergal J. et al. Ensembl 2023 Nucleic Acids Res 2023;51:D933–D941.

McLaren, William et al. The Ensembl Variant Effect Predictor Genome Biol 2016;17

Meng, Xiangchao et al. Hypoxia-inducible factor (HIF)-1alpha knockout accelerates intervertebral disc degeneration in mice Int J Clin Exp Pathol 2018;11

Mountjoy, Edward et al. An open approach to systematically prioritize causal variants and genes at all published human GWAS trait-associated loci Nat Genet 2021;53:1527–1533.

Mulder, Nicola, Opap, Kenneth Recent advances in predicting gene-disease associations F1000Res 2017;6

Pérez-Granado, Judith, Piñero, Janet, Furlong Laura I Benchmarking post-GWAS analysis tools in major depression: Challenges and implications Front Genet 2022;13

Piñero, Janet et al. The DisGeNET knowledge platform for disease genomics: 2019 update Nucleic Acids Res 2019;48:D845–D855.

Prokunina, Ludmila, Alarcón-Riquelme Marta E. Regulatory SNPs in complex diseases: Their identification and functional validation Expert Rev Mol Med 2004;6

Raudvere, Uku et al. g:Profiler: a web server for functional enrichment analysis and conversions of gene lists (2019 update) Nucleic Acids Res 2019;47:W191–W198.

Sollis, Elliot et al. The NHGRI-EBI GWAS Catalog: knowledgebase and deposition resource Nucleic Acids Res 2023;51:D977–D985.

Song, You Qiang et al. Lumbar disc degeneration is linked to a carbohydrate sulfotransferase 3 variant Journal of Clinical Investigation 2013;123:4909–4917.

Uffelmann, Emil et al. Genome-wide association studies Nature Reviews Methods Primers 2021;1

Williams, Frances M.K. et al. Novel genetic variants associated with lumbar disc degeneration in northern Europeans: A meta-analysis of 4600 subjects Ann Rheum Dis 2013;72:1141–1148.

Xu, Kang et al. Sp1 downregulates proinflammatory cytokine-induced catabolic gene expression in nucleus pulposus cells Mol Med Rep 2016;14:3961–3968.

